# The genome of *Litomosoides sigmodontis* illuminates the origins of Y chromosomes in filarial nematodes

**DOI:** 10.1101/2023.08.02.550553

**Authors:** Lewis Stevens, Manuela Kieninger, Brian Chan, Jonathan M D Wood, Pablo Gonzalez de la Rosa, Judith Allen, Mark Blaxter

**Affiliations:** Tree of Life, Wellcome Sanger Institute, Cambridge CB10 1SA, UK; Lydia Becker Institute of Immunology and Inflammation, Wellcome Centre for Cell-Matrix Research, Faculty of Biology, Medicine & Health, University of Manchester, Manchester M13 9PT, UK

**Author notes:** Correspondence should be addressed to Lewis Stevens or Mark Blaxter.

## Abstract

Heteromorphic sex chromosomes are usually thought to have originated from a pair of autosomes that acquired a sex-determining locus and subsequently stopped recombining, leading to degeneration of the sex-limited chromosome. In contrast, in rhabditid nematodes, sex is determined by an X-chromosome counting mechanism and males are hemizygous for one or more X chromosomes (XX/X0). Some species of filarial nematodes, including important parasites of humans, have heteromorphic XX/XY systems. It has been assumed that sex is determined by a Y-linked locus in these species. However, karyotypic analyses suggested that filarial Y chromosomes are derived from the unfused autosome following an X-to-autosome fusion. Here, we generated a chromosome-level reference genome for *Litomosoides sigmodontis*, a filarial nematode with the ancestral filarial karyotype and sex determination mechanism (XX/X0). We mapped the assembled chromosomes to the rhabditid nematode ancestral linkage (or Nigon) elements. We found that the *L. sigmodontis* X chromosome was formed from a fusion of NigonX (the ancestrally X-linked element) and NigonD (ancestrally autosomal) that occurred in the last common ancestor of all filarial nematodes. In the two filarial lineages with XY systems, the X chromosomes were formed from two recent and independent fusions of the ancestral X chromosome with different autosomal Nigon elements. In both lineages, the region shared by the neo-X and neo-Y chromosomes is within the ancestrally autosomal portion of the X, confirming that the filarial Y chromosomes are derived from unfused autosomes. Sex determination in XY filarial nematodes therefore likely continues to operate *via* the ancestral X-chromosome counting mechanism, rather than *via* a neo-Y-linked sex-determining locus.

## Introduction

Genetic sex determination is often associated with the presence of heteromorphic sex chromosomes, where males and females differ in their karyotypes. The presence of a distinct sex-limited chromosome often determines sex: Y in males of XY species, such as mammals and Diptera, and W in females of ZW species, such as birds and Lepidoptera (Bull 1983). The classical model of heteromorphic sex chromosome evolution posits that they originate when a pair of autosomes acquire a sex-determining locus and subsequently stop recombining (Rice 1996; Charlesworth 1996; Bergero & Charlesworth 2009). The lack of recombination in the sex-limited chromosomes (i.e. the Y or W) leads to degeneration. Sex-limited chromosomes tend to have fewer genes, accumulate repeats, and show extensive degeneration of coding loci, but are maintained because of the essential function of the sex determination loci they carry (Charlesworth 1996; Charlesworth & Charlesworth 2000; Bachtrog 2013). Sex chromosomes can evolve rapidly, including through fusion events between autosomes and X or Z chromosomes. In some lineages, neo-Y or neo-W chromosomes have arisen and are often derived from the now-haploid partner of a new autosome-X or -Z fusion (White 1973; McAllister & Charlesworth 1999; Carvalho & Clark 2005). The chromosomes recruited to these new Y or W roles often show ongoing degeneration in the same manner as much older Y or W chromosomes (Yi & Charlesworth 2000; Bachtrog & Charlesworth 2002; Carvalho & Clark 2005).

Many species have sex chromosomes but lack Y or W chromosomes. In these species, the heterogametic sex is haploid for the sex chromosome and sex is determined via a mechanism that assesses the autosome-to-sex chromosome ratio (Bridges 1921; Nigon 1951). The most common system is XX/X0 (X-null) sex determination, where males have only one X and produce sperm that either carry or do not carry an X. There is no male-determining locus and no pseudoautosomal region. The transition from XY or ZW to X0 or Z0 requires the evolution of a mechanism that assesses the autosome-to-X- or Z-chromosome ratio. With this in place, the Y or W is free to degenerate and be lost. In some lineages with X0 systems, *de novo* evolution of Y has been observed (e.g. in grasshoppers (White 1954)). As in XY and ZW species, the X chromosomes of X0 species are frequently involved in fusion events with autosomes (Gonzalez de la Rosa et al. 2021). In such cases, the unfused autosomal partner of the fused autosome could behave as a neo-Y chromosome, present only in males where it pairs (and thus shares a “pseudoautosomal region”) with the formerly autosomal part of the extended X. The neo-Y in these situations will be maintained if its fused homologue carries loci that are haplo-insufficient, until such time as haplo-insufficency and dosage compensation are resolved. Thus, neo-Y chromosomes in XX/X0 species may degenerate rapidly.

In the phylum Nematoda, the vast majority of species have an XX/X0 sex chromosome system. In *Caenorhabditis elegans*, where the mechanisms of sex determination have been well-studied, the number of X chromosomes is communicated by a set of X-linked loci termed X-signal elements (XSEs) (Gladden et al. 2007). The expression of the XSEs from the two copies of the X chromosome in *C. elegans* hermaphrodites (which are modified females capable of producing sperm) is sufficient to repress the activity of *xol-1*, the primary sex-determining gene. In contrast, XSE expression in males, which possess only a single X chromosome, is insufficient to suppress *xol-1*. Filarial nematodes (from family Onchocercidae within suborder Spirurina (De Ley & Blaxter 2004)), many of which are important parasites of humans, are unusual amongst nematodes in that several species have an XX/XY heteromorphic sex chromosome system. Filarial XY systems are proposed to have evolved twice independently from the ancestral XX/X0 system, once in the last common ancestor of *Dirofilaria* and *Onchocerca* and once in the last common ancestor of *Brugia* and *Wuchereria* (Post 2005; Lefoulon et al. 2015). It has often been assumed that sex determination in these species is driven by a sex-determining locus present on the Y, by analogy to XY systems in other taxa (e.g. (Wang et al. 2022)). However, karyotype analyses found that the evolution of XY systems in both filarial lineages coincided with an X:autosome fusion, suggesting that the Y chromosome may be derived from an unfused autosome (Post 2005).

The increasing availability of chromosome-level reference genomes has led to a renewed focus on chromosome evolution in recent years. Using the genomes of extant species, the linkage groups present in the last common ancestor have been inferred for many taxa, including *Drosophila* (Bhutkar et al. 2008), vertebrates (Simakov et al. 2020), and Lepidoptera (Wright et al. 2023). Seven ancestral linkage groups, called Nigon elements, are predicted to have been present in the last common ancestor of nematodes in the order Rhabditida (Tandonnet et al. 2019; Gonzalez de la Rosa et al. 2021). The chromosomes of extant rhabditid species can be derived from combinations of these elements. For example, the *C. elegans* X chromosome is derived from a fusion between NigonX (the ancestrally X-linked Nigon element) and NigonN (ancestrally autosomal) that occurred in the last common ancestor of all *Caenorhabditis* species (Gonzalez de la Rosa et al. 2021). A recent analysis attempted to reinfer Nigon elements to understand the evolution of the filarial nematode XY sex chromosome system, but erroneously inferred six ancestral rhabditid linkage groups instead of seven (Foster et al. 2020). In doing so, the origins of the filarial Y chromosomes were obscured. In addition, the conflicting hypotheses regarding the number of chromosomes in the last common rhabditid ancestor have led some authors to avoid using the Nigon terminology, despite a seven-Nigon rhabditid ancestor being consistent with their data (Yoshida et al. 2023).

To clarify the chromosomal biology of the filarial nematodes, we generated a chromosome-level reference genome for the rodent-parasitic species *Litomosoides sigmodontis*, which has a 5A, XX/X0 karyotype (Post 2005) that is likely to match that of the ancestor of the Onchocercidae (Gonzalez de la Rosa et al. 2021). We show that the *L. sigmondontis* X chromosome is the product of an ancient fusion event between NigonX and NigonD that occurred in or prior to the last common ancestor of all filarial nematodes. By analysing chromosomally complete genome sequences and Nigon element assignments within a phylogenetic context, we confirm that the two lineages with XY sex chromosome systems have undergone two recent and independent X-autosome fusions involving different autosomes. We show that the homologous region between the neo-X and neo-Y chromosomes corresponds to the ancestrally autosomal portion of the neo-X chromosome, confirming that the filarial neo-Y chromosomes are derived from unfused autosomes. We hypothesise that sex determination in filarial nematodes with XY sex chromosome systems operates similarly to the X-to-autosome ratio mechanism described in *C. elegans* and not by the presence of a sex-determining locus on the Y chromosome, and that retention of the Y because of haploinsufficiency of the fused homologue in males is an evolutionary intermediate to loss.

## Results

### *The* L. sigmodontis *X chromosome is a fusion of NigonX and NigonD*

We used a combination of high-coverage PacBio HiFi data derived from a single female and Hi-C data derived from two separate pools of adult males and females to generate a chromosome-level reference genome for *L. sigmodontis* (Figure 1A; Table 1). We scaffolded all of the 65.9 Mb primary nuclear genome assembly into six chromosome-sized scaffolds, consistent with the published *L. sigmodontis* karyotype (five autosomes plus the X chromosome) (Post 2005). All six chromosomal scaffolds were similar in size (10.3 - 11.7 Mb) and were capped by arrays of the nematode telomeric repeat ([TTAGGC]_n_) at both ends (Figure S1). The assembly is highly complete (96.0% using Benchmarking Using Single Copy Orthologues [BUSCOs] (Manni et al. 2021) with the Nematoda odb10 dataset) and has high base-level accuracy (QV of 52.9, corresponding to one error every 194 kb). We also generated a circular mitochondrial genome (spanning 13.7 kb) and a circular *w*Ls *Wolbachia* endosymbiont genome (spanning 1.05 Mb).

**Figure 1:**
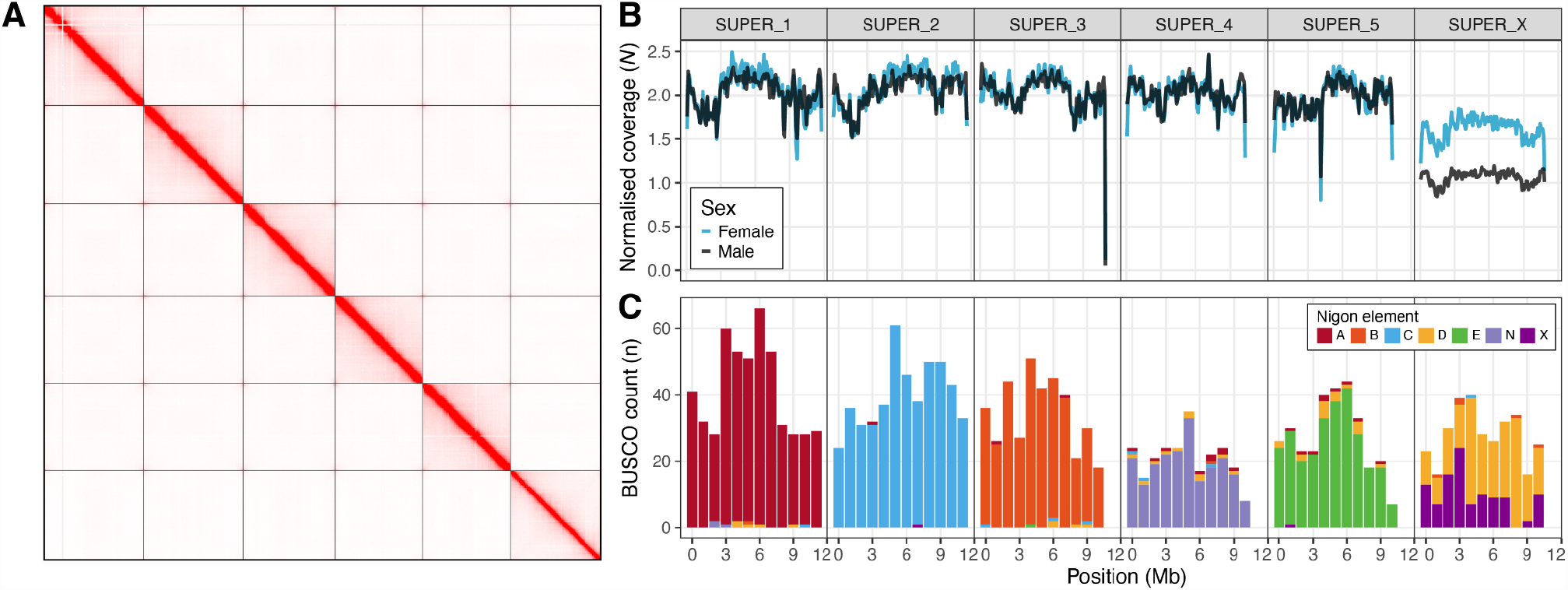
The *Litomosoides sigmodontis* X chromosome is a fusion of Nigon X and Nigon D. (A) Hi-C contact map for the nxLitSigm11.1 reference genome derived from the male Hi-C data. (B) Normalized male and female Hi-C read coverage in 100 kb windows in the six *L. sigmodontis* chromosomes. Normalised coverage (*N*) was calculated by dividing the coverage in each 100 kb window by the median autosomal coverage. (C) Distribution of counts of BUSCO genes in 500 kb windows in the six *L. sigmodontis* chromosomes coloured by their allocation to the seven Nigon elements (A-E, N, X).

**Table 1:** Metrics for chromosome-level reference genome of *Litomosoides sigmodontis* and three other filarial nematode species.

| **Accession information** |  |  |  |  |
| --- | --- | --- | --- | --- |
| Species | *Litomosoides sigmodontis* | *Dirofilaria immitis* | *Onchocerca volvulus* | *Brugia malayi* |
| BioProject | PRJEB64423 | - | PRJEB513 | PRJNA10729 |
| Reference | This work | (Gandasegui et al. 2023) | (Cotton et al. 2016) | (Foster et al. 2020) |
| **Reference genome metrics** |  |  |  |  |
| Span (Mb) | 65.9 | 89.4 | 96.4 | 88.2 |
| Scaffolds (n) | 6 | 27 | 708 | 197 |
| Scaffold N50 (Mb) | 10.9 | 15.2 | 25.5 | 14.2 |
| Span in chromosome-sized scaffolds (%) | 100.0 | 98.6 | 94.3 | 92.1 |
| Contigs (n)<sup>1</sup> | 28 | 167 | 1006 | 205 |
| Contig N50 (Mb)<sup>1</sup> | 5.0 | 4.2 | 1.0 | 10.3 |
| BUSCO completeness (%)<sup>2</sup> | 96.0 | 90.1 | 97.7 | 98.2 |
| BUSCO duplication (%)<sup>2</sup> | 0.4 | 0.5 | 0.9 | 0.6 |
| QV | 52.9 | - | - | - |
| **Gene set metrics** |  |  |  |  |
| Number of protein-coding genes | 8,660 | No gene set available | 12,109 | 10,878 |
| Number of transcripts | 9,551 | No gene set available | 13,436 | 15,985 |
| BUSCO completeness (%)<sup>2</sup> | 96.2 | No gene set available | 98.2 | 98.9 |

1. Contig values were calculated by splitting scaffolds at $\geq 10$ consecutive Ns.
2. Genome and gene set completeness was assessed using BUSCO (version 5.2.2) with the nematoda\_odb10 dataset (using the Augustus option when assessing genome completeness).

To identify the *L. sigmodontis* X chromosome, we aligned male and female Hi-C data to the genome and calculated per-base read coverage. Consistent with the reported karyotype, we found five scaffolds that had diploid coverage in both males and females and are thus identified as autosomes (Figure 1B). The remaining scaffold had haploid coverage in males and was identified as the X chromosome (Figure 1B). In females, this scaffold had approximately 80% of the coverage of the autosomal scaffolds (Figure 1B). We interpret this as being due to the fact that adult female *L. sigmodontis* carry large numbers of both male and female embryos retained *in utero*, and this results in sub-diploid coverage of the X.

We painted the autosomes and X chromosome using conserved BUSCO genes allocated to the seven Nigon elements (Gonzalez de la Rosa et al. 2021). The five autosomes were primarily composed of loci from a single Nigon element (NigonA, B, C, E, and N), suggesting that these chromosomes have not undergone fusion or fission since the last common rhabditid ancestor (Figure 1C). In contrast, the X chromosome was composed of genes from both NigonX and NigonD (Figure 1C). The NigonX and NigonD genes were highly intermixed, suggesting that sufficient evolutionary time has passed since the fusion event for intra-chromosomal rearrangements to have blended the distinct Nigon domains.

### Filarial nematodes with XY sex chromosomes have undergone recent X-autosome fusions

To understand the evolutionary origins of the filarial XY systems, we analysed previously reported karyotypes and Nigon assignments in three XY species with chromosome-level reference genomes (*Onchocerca volvulus* (Cotton et al. 2016), *Brugia malayi* (Foster et al. 2020), and *Dirofilaria immitis* (Gandasegui et al. 2023)) within the phylogenetic context of the Onchocercidae (Lefoulon et al. 2015) (Figure 2). Post (2005) previously inferred that the 5A, X0 karyotype observed in *L. sigmodontis, Acanthocheilonema viteae, Loa loa*, and *Setaria labiatopapillosa* was likely to be the ancestral filarial karyotype (Figure 2A). The NigonX+NigonD fusion therefore occurred in (or prior to) the last common filarial ancestor, which is consistent with the degree of intermixing between the NigonX and NigonD genes observed in the *L. sigmodontis* X chromosome.

**Figure 2:**
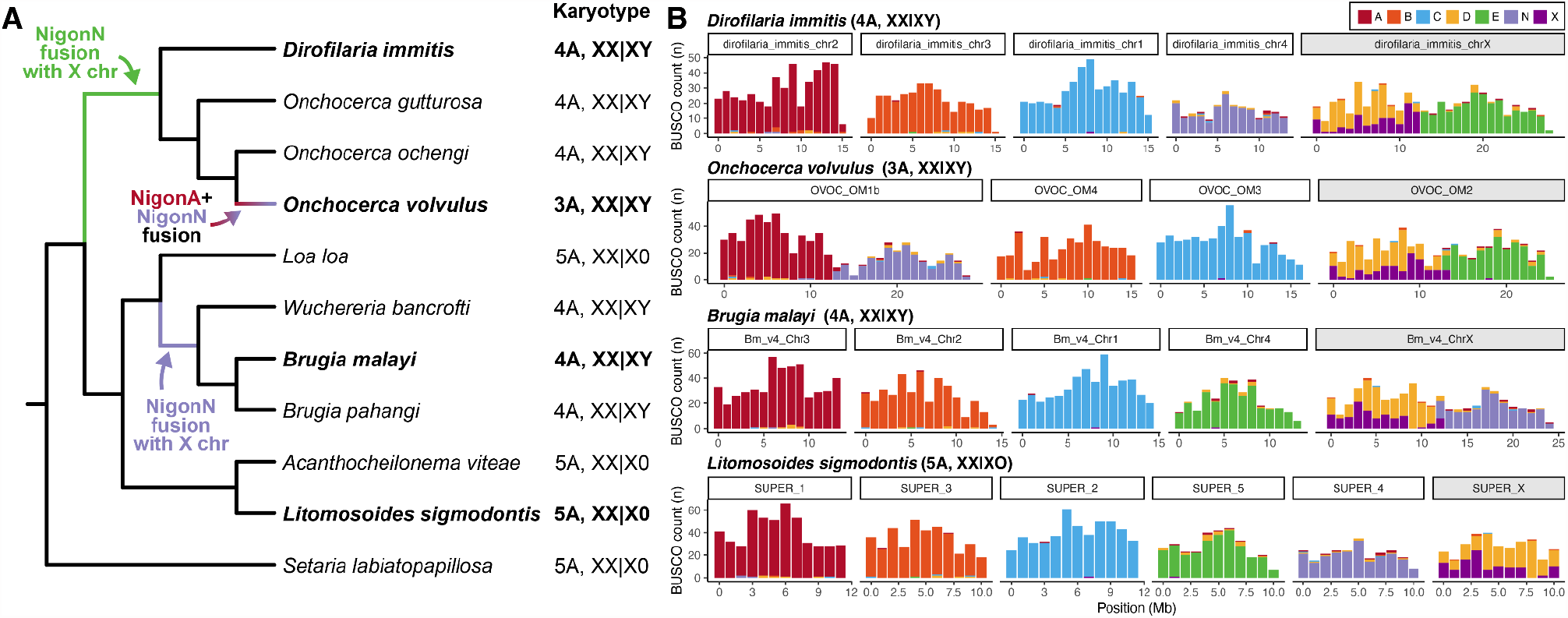
Filarial nematodes with XY sex chromosomes have undergone recent X:autosome fusions. (A) Cladogram of filarial nematodes derived from a supermatrix comprising 1,757 single-copy orthologues present in all 16 species. Reported karyotypes are indicated (Triantaphyllou 1983; Post 2005). Species with chromosome-level reference genomes figured in (B) are highlighted in bold. (B) Distribution of BUSCO genes coloured by their allocation to the seven ancestral Nigon elements in the chromosome-level assemblies of *Dirofilaria immitis, Onchocerca volvulus, Brugia malayi*, and *L. sigmodontis*. The scaffold that comprises the X chromosome is indicated in grey. Species names and karyotypes are indicated above each plot.

XY sex chromosome systems have evolved twice independently from X0 systems in filarial nematodes (Post 2005) (Figure 2A). In both cases, XY evolution coincided with an X-to-autosome fusion involving two different autosomes (Gonzalez de la Rosa et al. 2021) (Figure 2A). In the ancestor of *Dirofilaria* and *Onchocerca*, a fusion between NigonE and the X chromosome (NigonX+NigonD) led to a 4A, XY karyotype in *Dirofilaria immitis* (Figure 2B). Consistent with this, the *D. immitis* X chromosome (28.2 Mb) is approximately twice the size of the autosomes (which range in size from 14.0 to 15.6 Mb). An additional autosome-to-autosome fusion (between NigonA and NigonN) occurred in the lineage leading to *O. volvulus*, generating its 3A, XY karyotype (Figure 2B). A similar autosome-to-autosome fusion is likely to be present in *Onchocerca gibsoni* (Post 2005), which is closely related to *O. volvulus* (Lefoulon et al. 2015). Other *Onchocerca* species have 4A, XY karyotypes and therefore presumably lack this fusion (Post 2005). The X chromosomes of both *D. immitis* and *O. volvulus* retain a structure reflecting their origins. The NigonE partition remains distinct from the intermixed NigonX+NigonD and the fusion breakpoint retains features of nematode chromosome ends, such as higher repeat content (Figure 2B; Figure S2). We note that the *D. immitis* genome was scaffolded on the *O. volvulus* assembly (Gandasegui et al. 2023), and thus that there may be reference bias in the proposed chromosome structure.

In the ancestor of *Brugia* and *Wuchereria*, an independent fusion occurred between NigonN and the X chromosome, generating the 4A, XY karyotype seen in all species in this clade (Figure 2A) (Post 2005; Lefoulon et al. 2015). The fusion occurred at the opposite end of the X chromosome relative to the *Onchocerca* (and *Dirofilaria*) fusion (Figure S3-5). The X chromosome is again substantially larger than the autosomes (24.9 Mb versus 13.4 - 14.7 Mb) and the fusion strongly patterns the new chromosome. The NigonN partition remains distinct from the NigonX+NigonD partition and the fusion breakpoint is evident in the repeat distribution (Figure 2B; Figure S2). In summary, filarial XY chromosome systems have evolved twice, independently, from an ancestral 5A, X0 karyotype via distinct X-to-autosome fusion events.

### Filarial Y chromosomes are derived from unfused autosomes

Filarial Y chromosomes are hypothesised to be derived from unfused autosomes (Post 2005; Gonzalez de la Rosa et al. 2021). Under this model, the ancestrally autosomal portion of the X chromosome (derived from NigonE in *D. immitis* and *O. volulvus* and NigonN in *B. malayi*) should be diploid in males, forming a “pseudoautosomal region” (PAR), while the ancestrally X-derived portion (NigonX+NigonD) should remain haploid. To determine if this is the case, we aligned male- and female-specific short-read data to the *D. immitis, O. volvulus*, and *B. malayi* reference genomes and calculated per-base read coverage (Figure 3).

**Figure 3:**
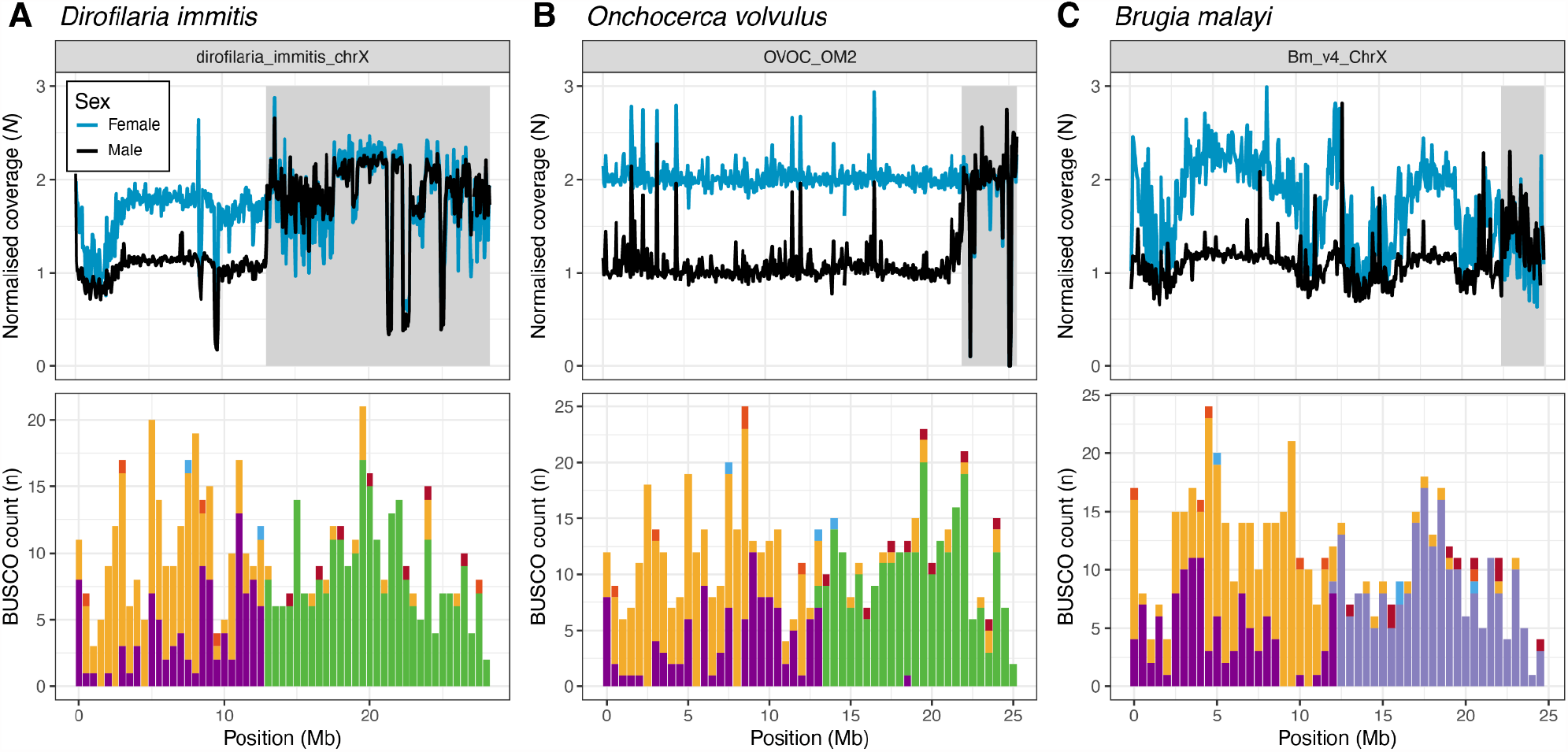
Filarial Y chromosomes are derived from unfused autosomes. Male and female read coverage and Nigon element partitions in the X chromosomes of (A) *D. immitis*, (B) *O. volvulus*, and (C) *B. malayi*. Normalised coverages (*N*) are calculated by dividing the coverage in each 50 kb window by the median autosomal coverage. The histogram of locations of BUSCO loci allocated to Nigon elements (coloured as in Figure 1C) are binned in 500 kb windows. Pseudoautosomal regions are shown by grey shading.

In *D. immitis*, the entirety of the NigonE partition of the X chromosome is diploid in males, forming a 15.2 Mb PAR, while the ancestrally X-derived partition is haploid (Figure 3A). The divergence between the X and Y chromosomes within the PAR is low, with an average of 1 SNP every 2,215 bp (Figure S6; Table S1). The *D. immitis* Y chromosome therefore appears to be an intact, unfused NigonE autosome. In *O. volvulus*, the pseudoautosomal region is substantially shorter and covers only 3.4 Mb of the NigonE partition of the X chromosome (Figure 3B). The remaining 8.6 Mb of the NigonE-derived sequence has haploid coverage in males, suggesting that the Y chromosome copy of the NigonE element has largely degenerated in *O. volvulus*. Based on analyses of the divergence between X and Y in other XY systems, the divergence between X and Y is likely to have been through several “strata” of regions which reflect the progressive but episodic loss of recombination along the former homologues. Consistent with this, SNP density is elevated in *O. volvulus* males in the region neighbouring the pseudoautosomal region boundary (Figure S6) and there is an average of 1 SNP every 314 bp within the PAR (Table S1). In addition, many of the *O. volvulus* Y contigs identified by (Cotton et al. 2016) align to the left boundary of the PAR, and show on average 13% nucleotide divergence (Figure S7). In *B. malayi*, there is a 2.6 Mb PAR, which is found in the NigonN partition of the chromosome (Figure 3C). As in *O. volvulus*, a large portion of the NigonN partition is haploid in males, suggesting that the *B. malayi* Y chromosome has also undergone degeneration. Heterozygous SNP density is also elevated in the region neighbouring the PAR boundary in males (Figure S6) and many of the previously identified Y chromosome contigs (Foster et al. 2020) align to the left of the PAR boundary (Figure S7). In summary, the PARs in all three species occur within the ancestrally autosomal portions of the X chromosomes, confirming that filarial Y chromosomes are derived from the unfused autosome following X-to-autosome fusion.

## Discussion

We have assembled a chromosomally complete reference genome for the filarial nematode *L. sigmodontis* and confirmed that it has a karyotype of five autosomes and an X chromosome (Post 2005), that the origin of these chromosomes can be explained by the Nigon element model of seven ancestral linkage groups in Rhabditida (Gonzalez de la Rosa et al. 2021), and that the X chromosome is a relatively old fusion of the NigonX and NigonD elements. We combined these new data with publicly available data for other filarial species to explore the origins of the XX/XY sex chromosome system of two clades of filarial nematodes that include major parasites of humans (*Onchocerca* plus *Dirofilaria*, and *Brugia* plus *Wuchereria*). We confirm the independent origin of these XX/XY systems through the fusion of distinct autosomes to the NigonX+NigonD ancestral X chromosome and show that the pseudoautosomal region (PAR) shared by the neo-X and neo-Y chromosomes is found within the ancestrally autosomal portion in three species with chromosome-level reference genomes.

Degeneration of sex-limited chromosomes (Y or W) is usually ascribed to the suppression of recombination around a sex-determining locus that progressively spreads across the chromosome (Bergero & Charlesworth 2009). In *B. malayi* and *O. volvulus* the Y is degenerated, and the PARs comprise a small, telomere-adjacent segment comprising a quarter to a fifth of the ancestrally autosomal partition in the neo-X chromosome. However, the PARs of these two species are not homologous, as they derive from distinct Nigon elements. If they contain a male-determining locus, these loci must, like the fusions, have independent origins.

The X chromosome of nematodes (marked by the loci defining NigonX) is frequently involved in fusion events with autosomes (Gonzalez de la Rosa et al. 2021). This means that many nematode species have XX/X0 sex determination based on fused X chromosomes, and thus that in the context of an XX/X0 system, X-autosome fusions are likely to be resolved by eventual loss of the neo-Y and retention of the XX/X0 mechanism. On this basis, we suggest that sex determination in the filarial XX/XY species is still likely to be functionally mediated *via* an X:A ratio counting mechanism, and that the neo-Y chromosomes play no genetic part in sex determination. Like filarial XX/XY species, the free-living nematode *Pristionchus exspectatus* has a neo-XY system that originated through a very recent fusion of an ancestral X chromosome (which was solely NigonX) with an autosome (NigonN) (Yoshida et al. 2023). Female *P. exspectatus* carry two copies of the fused NigonX+NigonN X chromosome and the male has one NigonX+NigonN X and one NigonN chromosome (the Y). As in filarial nematodes, there are no data indicating that the NigonN neo-Y has any functional role in sex determination. Instead, we suggest that the neo-Y chromosomes in filarial nematodes and in *P. exspectatus* are necessary intermediates in the amelioration of the newly X-linked, ancestrally-autosomal chromosome portion to haploidy. This amelioration likely includes the spread of the X-chromosome dosage compensation system over the newly sex-linked compartment and the resolution of any haploinsufficient loci (such as masked deleterious alleles) in the ancestrally autosomal portion of the X chromosome. This model suggests that all lineages where the X chromosome has fused to an autosome, including *Caenorhabditis* (Gonzalez de la Rosa et al. 2021), once possessed and have lost a neo-Y chromosome. In this context, we note that B chromosomes have been observed in *Onchocerca* species (Procunier & Hirai 1986; Post et al. 1989), which could be fragments of a degenerating Y.

Why have the Y chromosomes of *O. volvulus* and *B. malayi* degenerated in the absence of sex-determining loci? The pattern of degeneration of the neo-Y chromosomes in *B. malayi* and *O. volvulus* is striking but may be the result of the patterning of recombination in the neo-X chromosome. In a *C. elegans* strain that has undergone an X-to-autosome fusion, generating a neo-XY system, the landscape of recombination between the neo-X and Y chromosomes was repatterned such that crossover events were concentrated into a short region of the ancestrally autosomal segment distal to the fusion point (Henzel et al. 2011). If recombination was repatterned in a similar way after the X-to-autosome fusions in the ancestors of *B. malayi* and *O. volvulus*, the relative lack of recombination in the region proximal to the fusion point could drive the observed ratchet-like and stepwise pattern of Y chromosome degeneration spreading from the point of fusion.

It is striking that, in contrast to *O. volvulus* and *B. malayi*, the neo-Y in *D. immitis* shows very little degeneration and retains alignable coverage (the PAR) along the full length of its homologue fused to the X chromosome. While this fusion appears to be homologous to the one in *O. volvulus*, under this assumption, the lack of degeneration of the *D. immitis* Y is problematic: why is it not similarly affected by degeneration? It seems unlikely that recombination repatterning or the evolution of dosage compensation would follow distinct evolutionary paths in the two lineages. One explanation could be that the *Dirofilaria* and *Onchocerca* fusions are in fact independent, and the *D. immitis* Y chromosome is less degenerated because the fusion was more recent. This requires two independent fusions of the ancestral X and NigonE in the same orientation, which is possible but seems unlikely. Alternatively, we note that the contig-level *D. immitis* genome (Gomes-de-Sá et al. 2022) was scaffolded on the *O. volvulus* chromosomal assembly (Gandasegui et al. 2023), and there is a possibility that this reference-guided approach may have categorically biased the outcome. Without independent chromosomal assembly, characterising the recombination landscape in *D. immitis*, or generating chromosome-level reference genomes for other species in this clade, we are not able to distinguish between these possibilities.

A recent analysis assumed that the ancestral nematode had an XX/XY sex-determination system, that the XX/X0 system observed in the majority of species is derived (Wang et al. 2022), and that existing nematode XX/XY species retain the ancestral mechanism. This model requires a very large number of independent and genetically homoplastic origins of an XX/X0 system across the phylum. It relies on questionable identification of “Y-like” chromosomes in some *Trichuris* species (Spakulová & Casanova 2004; Spakulová et al. 1994) which could equally have recent origins similar to those defined for filarial nematodes (this work) and for *P. exspectatus* (Yoshida et al. 2023). We note that B chromosomes have been described in *Trichuris* species (Goswami 1978) as in the XY filaria. There is no compelling evidence for any loci that might have roles in sex determination that might have travelled with a ghost Y chromosome, and indeed the neo-Y chromosomes of the four fully-sequenced XY nematode species have three distinct Nigon element origins. The evolution of neo-Y chromosomes is common and often involves X-to-autosome fusion, for example in *Drosophila* (Bachtrog 2013; Dupim et al. 2018).

Nematodes show great variability in karyotype, genome organisation, and reproductive modes (Mitreva et al. 2005; Denver et al. 2011; Gonzalez de la Rosa et al. 2021). Importantly this diversity is present within species-rich clades and many transitions have multiple independent occurrences, such as the homoplastic origins of the XX/XY system in filarial nematodes. Coupled with a wealth of new chromosomally-complete genomes, the emerging synthesis of patterns of chromosomal evolution in the phylum promises to become a rich ground to explore larger questions of the drivers of and constraints on evolution. The ancestral linkage groups defined for Rhabditida, the Nigon elements (Gonzalez de la Rosa et al. 2021), are key organising principles in this endeavour.

## Methods

### Collection of adult *L. sigmodontis*

The life cycle of *L. sigmodontis* was maintained at the University of Manchester in accordance with the United Kingdom Animals (Scientific Procedures) Act of 1986 under Project License PP4115856. Mongolian jirds (*Meriones unguiculatus*) were infected with 80 infective L3 larvae by intraperitoneal injection; animals were maintained at 27°C and 75% relative humidity. Euthanasia was performed by cervical dislocation 120 days post-infection and adult worms were collected from the peritoneal cavity into PBS. Individual worms of each sex were placed in Fluidx tubes chilled on dry ice. Two pools comprising 10 adult males and 10 adult females were collected in the same way. The samples were stored at -80°C.

### DNA extraction and genome sequencing

We extracted DNA from a whole adult female *L. sigmodontis* using the New England Biolabs Monarch^®^ HMW DNA Extraction Kit for Tissue using the manufacturers’ standard input protocol with the following modifications. Briefly, we stored the tissue at -70°C and transferred it to a 1.5 ml Monarch pestle tube before disruption. We digested the tissue for 30 min at 56°C with 600 rpm before adding RNase A. We eluted the DNA in 200 μl Monarch^®^ gDNA Elution Buffer II. The DNA was sheared twice with the Megaruptor^®^ 3 from Diagenode with a velocity setting of 30 and 31. The sheared DNA was SPRI cleaned with the ProNex® Size-Selective Purification System in a ratio of 1x (v/v) and eluted in 48 μl PacBio Elution buffer. The average DNA fragment size was ∼15.8 kb.

A PacBio library was prepared by the Long Read Team of the Scientific Operations core at the Wellcome Sanger Institute using the PacBio Low DNA Input Library Preparation Using SMRTbell Express Template Prep Kit 2.0. The library was sequenced on a single PacBio Sequel IIe flow cell.

Hi-C library preparation and sequencing were performed by the Long Read Team of the Scientific Operations core at the Wellcome Sanger Institute. Two pellets comprising 10 adult males and 10 adult females were processed using the Arima Hi-C version 2 kit following the manufacturer’s instructions. Illumina libraries were prepared using the NEBNext Ultra II DNA Library Prep Kit. Each library was sequenced on one-eighth of a NovaSeq S4 lane using paired-end 150 bp sequencing.

### Genome assembly

We removed adapter sequences from the PacBio HiFi data using HiFiAdapterFilt (Sim et al. 2022). We used Jellyfish 2.3.0 (Marçais & Kingsford 2011) to count *k*-mers of length 31 in each read set and GenomeScope 2.0 (Ranallo-Benavidez et al. 2020) to estimate genome size and heterozygosity. We first generated a preliminary assembly of the PacBio HiFi data using hifiasm v0.16.1-r375 (Cheng et al. 2021). We randomly subsampled 10% of the male Hi-C reads using samtools 1.14 (Danecek et al. 2021) and aligned them to the hifiasm primary assembly using bwa mem 0.7.17-r1188 (Li 2013), filtered out PCR duplicates using piccard 2.27.1-0 (available at http://broadinstitute.github.io/picard/), and scaffolded the assembly using YaHS (Zhou et al. 2023). We ran BlobToolKit 2.6.5 (Challis et al. 2020) on the scaffolded assembly and used the interactive web viewer to manually screen for scaffolds derived from non-target organisms. We identified a single scaffold that was of non-nematode origin, which corresponded to the *Wolbachia* endosymbiont genome, which we processed separately (see below). After removing the *Wolbachia* scaffold, we used MitoHiFi 2.2 (Uliano-Silva et al. 2022) to extract and annotate the mitochondrial genome. Finally, we removed residual haplotypic duplication from each assembly using purge_dups 1.2.5 (Guan et al. 2020) and used seqkit (Shen et al. 2016) and BUSCO 5.2.2 (Manni et al. 2021) with the nematoda_odb10 lineage to calculate assembly metrics and assess biological completeness. We assessed base-level accuracy and *k*-mer completeness using Merqury 1.3 (Rhie et al. 2020).

### Hi-C scaffolding and manual curation

To generate a chromosome-level reference genome for *L. sigmodontis*, we scaffolded the primary assembly using the male Hi-C data as previously described. We manually curated the scaffolded assembly as in (Chow et al. 2016) and used PretextView 0.2.5 (available at https://github.com/wtsi-hpag/PretextView) to manually inspect the Hi-C signal. Two contigs were identified as haplotypic duplication and were removed from the primary assembly. A final Hi-C contact map was generated using Juicer 2.0 (Durand et al. 2016) and Juicebox 1.11.08 (available at https://github.com/aidenlab/Juicebox) and is shown in Figure 1A.

### *Wolbachia* genome assembly

The *Wolbachia* contig produced by hifiasm was not circular. To generate a circular *Wolbachia* genome, we aligned the PacBio HiFi reads to the hifiasm contig using minimap2 2.24-r1122 (Li 2018) and extracted mapped reads. We assembled the mapped reads using flye 2.9-b1768 (Kolmogorov et al. 2018), which yielded a circular 1.05 Mb contig that shared 100% nucleotide identity with the publicly available circular *w*Ls genome (GCA_013365435.1) (Lefoulon et al. 2020) and differed in length by just 1 bp. We annotated the genome using Prokka 1.14.6 (Seemann 2014) and rotated the genome to start with HemE, as in (Vancaester & Blaxter 2023).

### Gene prediction

Prior to predicting protein-coding genes, we used Earl Grey 2.0 (Baril et al. 2022) to perform *de novo* repeat identification on the *L. sigmodontis* genome. We provided the curated repeat library to RepeatMasker 4.1.2-p1 (Smit et al. 2015) to soft-mask transposable elements. As we did not have access to RNA-seq data for *L. sigmodontis*, we used BRAKER2 2.1.6 (Brůna et al. 2021), which uses protein alignments from related species as evidence. To create a protein database, we ran BUSCO (using the nematoda_odb10 lineages) on the *L. sigmodontis* nxLitSigm1.1 reference genome and reference genomes for ten other filarial nematode species (Table S2). We used busco2fasta (available at https://github.com/lstevens17/busco2fasta) to identify and extract the protein sequences of BUSCO genes that were present and single-copy in at least four of the 11 species. The resulting protein database, comprising 32,050 protein sequences, was provided to BRAKER2 to predict genes in the soft-masked nxLitSigm11.1 reference genome. The number of predicted genes (8,660) is lower than reported for *Brugia malayi* and *Onchocerca volvulus* (Table 1) and is likely an underestimate (owing, in part, due to the lack of RNA-seq data).

### Coverage estimation in *L. sigmodontis*

To estimate per-base read coverage of each chromosome in the *L. sigmodontis* reference genome, we randomly subsampled 10% of the male and female Hi-C reads using samtools and aligned them to the genome using bwa mem, as previously described. We used BEDtools v2.30.0 (Quinlan & Hall 2010) to generate non-overlapping windows of various sizes and mosdepth 0.3.3 (Pedersen & Quinlan 2018) to calculate coverage in each window.

### Nigon painting

To analyse the distribution of Nigon elements in filarial chromosomes, we ran BUSCO using the nematoda odb10 dataset on chromosome-level reference genomes for *L. sigmodontis, D. immitis* (Gandasegui et al. 2023), *O. volvulus* (Cotton et al. 2016), and *B. malayi* (Tracey et al. 2020) (see Table S2 for accessions). We used buscopainter (available at https://github.com/lstevens17/buscopainter) to assign each BUSCO gene to a Nigon element, as defined by (Gonzalez de la Rosa et al. 2021).

### Filarial phylogeny

To infer a filarial phylogeny, we used busco2fasta (available at https://github.com/lstevens17/busco2fasta) to identify and extract the protein sequences of 1,757 BUSCO genes that were single-copy in the nxLitSigm11.1 reference genomes and genomes of ten other filarial species (Table S2). We aligned the protein sequences using FSA 1.15.9 (Bradley et al. 2009) and concatenated the alignments of each single-copy orthogroup into a supermatrix using catfasta2phyml v1.1.0 (available at https://github.com/nylander/catfasta2phyml). We used IQ-TREE 2.2.0.3 (Minh et al. 2020) to infer the filarial phylogeny under the LG substitution model (Le & Gascuel 2008) with gamma-distributed rate variation among sites. We visualised the resulting species tree using the iTOL web server (Letunic & Bork 2016).

### Repeat distribution

To analyse repeat distribution across each chromosome, we used Red 2.0 (Girgis 2015) with a *k-*mer length of 13 to identify repetitive sequences in the *L. sigmodontis, D. immitis, O. volvulus*, and, *B. malayi* genomes. We calculated repeat density in non-overlapping 200 kb windows using BEDtools v2.30.0. We note that the repetitive proportion estimated for *L. sigmodontis* by Red (44%) is substantially higher than estimated using earlGrey (6%), which is likely due to earlGrey being designed to identify full-length transposable elements, whereas Red identifies repetitive sequences of any size, the majority of which are not transposable elements.

### Inferring the PAR coordinates in filarial species with XY chromosomes

To define pseudoautosomal regions in the X chromosomes of *D. immitis, O. volvulus*, and *B. malayi*, we mapped a set of male- and female-derived paired-end short-read Illumina reads (see Table S3 for accessions) to each genome using bwa mem and calculated coverage in non-overlapping 100 kb windows, as previously described. We used a similar definition to (Foster et al. 2020) to identify pseudoautosomal regions as large (> 1 Mb in length), semi-contiguous regions of the X chromosome where no bins were female-dominated (male-to-female coverage ratio < 0.8). To do this, we normalised male and female read coverage by dividing the coverage in each 100 kb window by the autosomal median coverage (the median of all 100 kb windows excluding those on the X chromosomes). We then merged all non-female-dominated bins that were separated by 200 kb or less using BEDtools and retained only windows that were at least 1 Mb in size. Using this approach, we identified a single PAR in *B. malayi* (Bm_v4_ChrX:22,300,000-24,943,668) and in *O. volvulus* (OVOC_OM2:22,100,000-25,485,961).

In *D. immitis*, there were two non-overlapping windows larger than 1 Mb, but one of these was a 1.4 Mb region that showed low coverage in both males and females (dirofilaria_immitis_chrX:700,000-2,100,000), whereas the other was a 15.2 Mb region that had diploid coverage in both males and females (dirofilaria_immitis_chrX:13,000,000-28,232,375) (Figure 3A).

### Estimating divergence between X and Y chromosomes using variant calling

To estimate the divergence between the X and Y chromosomes of *D. immitis, O. volvulus*, and *B. malayi*, we used the BAM files generated previously to call variants in each read set using DeepVariant 1.4.0 (Poplin et al. 2018). We filtered out any variant labelled with ‘RefCall’ and used BCFtools 1.15 (Danecek et al. 2021) to filter the resulting VCF files to contain only heterozygous biallelic single nucleotide polymorphisms (SNPs). For each read set, we calculated heterozygous SNP density across the genome using 50 kb, non-overlapping windows using BEDtools. To plot the average divergence between the X and Y chromosomes, we inferred the sex of each individual based on the coverage of the X chromosome (Table S3) and calculated the mean SNP density in each 50 kb window for all male read sets using BEDtools. We also calculated the average SNP density for each male across the entirety of the PARs inferred previously (Table S1).

### Comparing divergence between Y contigs

We removed the 63 *B. malayi* Y contigs identified by (Foster et al. 2020) and 148 *O. volvulus* Y contig identified by (Cotton et al. 2016) from each reference genome and then aligned them to the remaining contigs/scaffolds using NUCmer, which is part of the MUMmer package 3.1 (Kurtz et al. 2004). We retained only one-to-one alignments (using delta-filter, which is also part of the MUMmer package) that were 1 kb or longer. We calculated alignment coverage in non-overlapping 50 kb windows using BEDtools, as previously described. We calculated the mean alignment identity by multiplying the identity of each alignment by its length and dividing by the total length of all alignments.

## Data availability

The *L. sigmodontis* PacBio HiFi data are available at ENA under the accession ERR11560319. The male and female *L. sigmodontis* Hi-C data are available under the accessions ERR11560320 and ERR11560321, respectively. The nxLitSigm11.1 reference genome is available under the BioProject PRJEB64423. The alternate haplotype assembly is available under the BioProject PRJEB64424. The *Wolbachia* genome is available under the accession PRJEB64425. Accessions for publicly available reference genomes are in Table S2. Accessions for publicly available short-read data are available in Table S3. Code and intermediate files associated with the analyses and figures are available at https://github.com/lstevens17/litomosoides_MS.

## Acknowledgements

This work was supported by Wellcome Trust Grants 206194 and 218328. For the purpose of Open Access, the author has applied a CC BY public copyright licence to any Author Accepted Manuscript version arising from this submission. *L. sigmodontis* maintenance at the University of Manchester was funded by the Medical Research Council UK (MR/K01207X/1) and Wellcome Trust (106898/A/15/Z). We thank the Sanger Scientific Operations Long Read team for sequencing and the Genome Reference Informatics Team in Tree of Life for genome curation assistance. We thank members of team301 and Kamil Jaron for feedback on the manuscript and Emmelien Vancaester for assistance with assembling and annotating the *Wolbachia* genome.

