## Supplementary figures for "The genome of *Litomosoides sigmodontis* illuminates the origins of Y chromosomes in filarial nematodes"

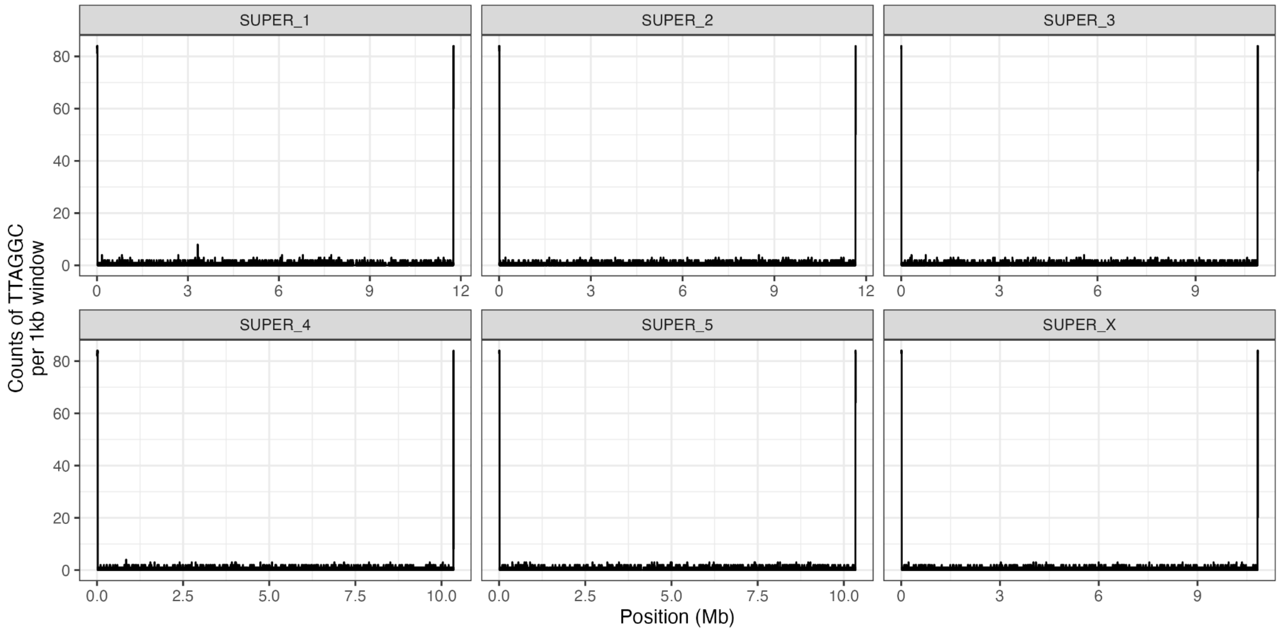


**Figure S1: Telomeric repeat sequence in *Litomosoides sigmodontis* nxLitSigm11.1 reference genome**
Counts of the nematode telomeric repeat sequence (TTAGGC) in 1 kb windows in the nxLitSigm11.1 reference genome.

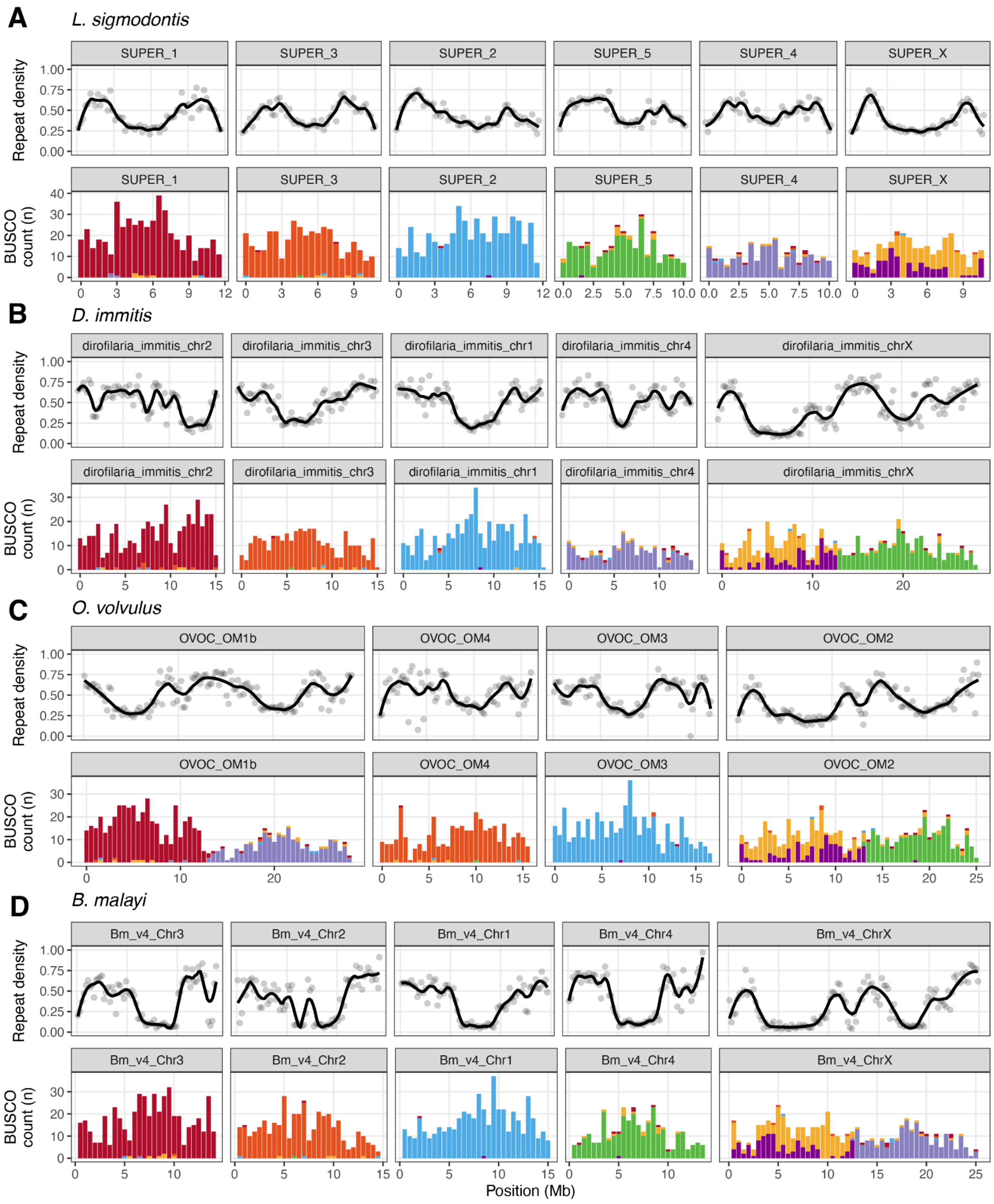


**Figure S2: Repeat distributions in the genomes of four filarial nematode species**

Repeat and Nigon element distributions in the genomes of (A) *L. sigmodontis*, (B) *D. immitis*, (C) *O. volvulus*, and (D) *B. malayi*. Repetitive sequences were identified using Red with a kmer length of 13 and repeat densities were calculated in 200 kb, non-overlapping windows. Lines represent LOESS smoothing functions fitted to the data. Distribution of counts of BUSCO genes in 500 kb windows in the six *L. sigmodontis* chromosomes by their allocation to the seven Nigon elements (coloured as in Figure 1C). The repetitive proportion estimated for *L. sigmodontis* genome by Red (44%) is substantially higher than estimated using earlGrey (6%), which is likely due to earlGrey being designed to identify full-length transposable elements, whereas Red identifies repetitive sequences of any size, the majority of which are not transposable elements.

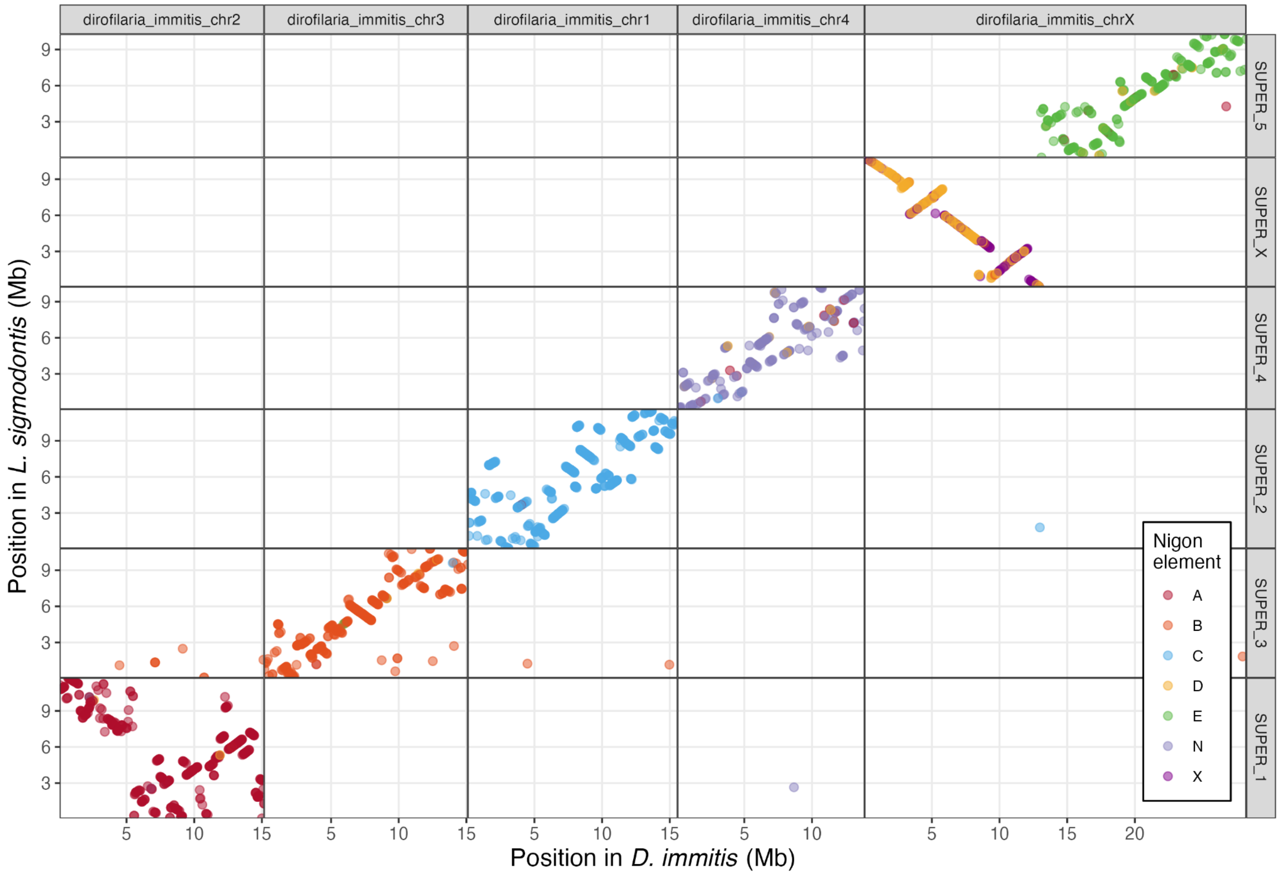


**Figure S3: Synteny between *D. immitis* and *L. sigmodontis***

The relative position of 1,979 BUSCO genes in the *D. immitis* and *L. sigmodontis* genomes. BUSCO genes are coloured by their Nigon assignment. Unassigned BUSCO genes are not shown. *D. immitis* chromosomes are ordered as in Figure 2B.

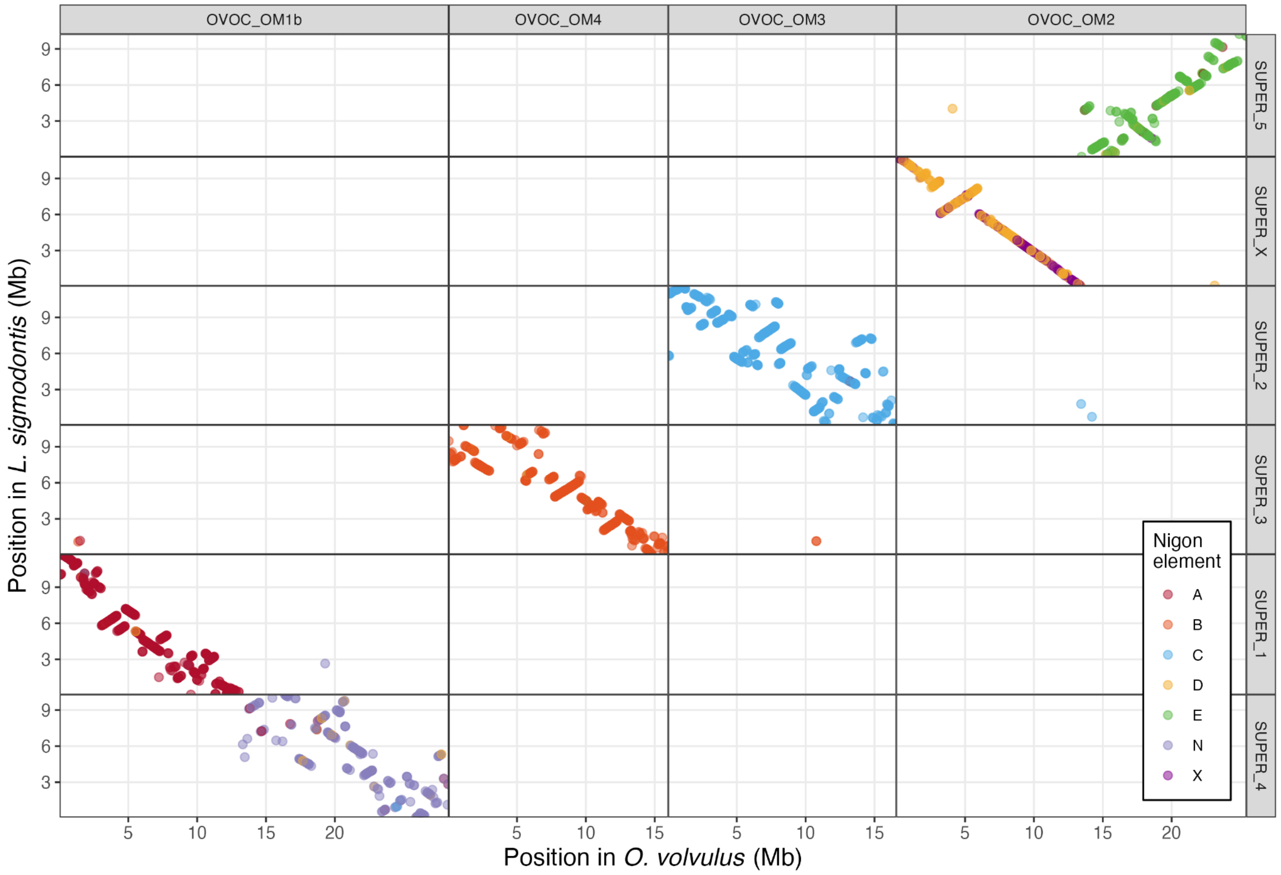


**Figure S4: Synteny between *O. volvulus* and *L. sigmodontis***
The relative position of 2,198 BUSCO genes in the *O. volvulus* and *L. sigmodontis* genomes. BUSCO genes are coloured by their Nigon assignment. Unassigned BUSCO genes are not shown. *O. volvulus* chromosomes are ordered as in Figure 2B.

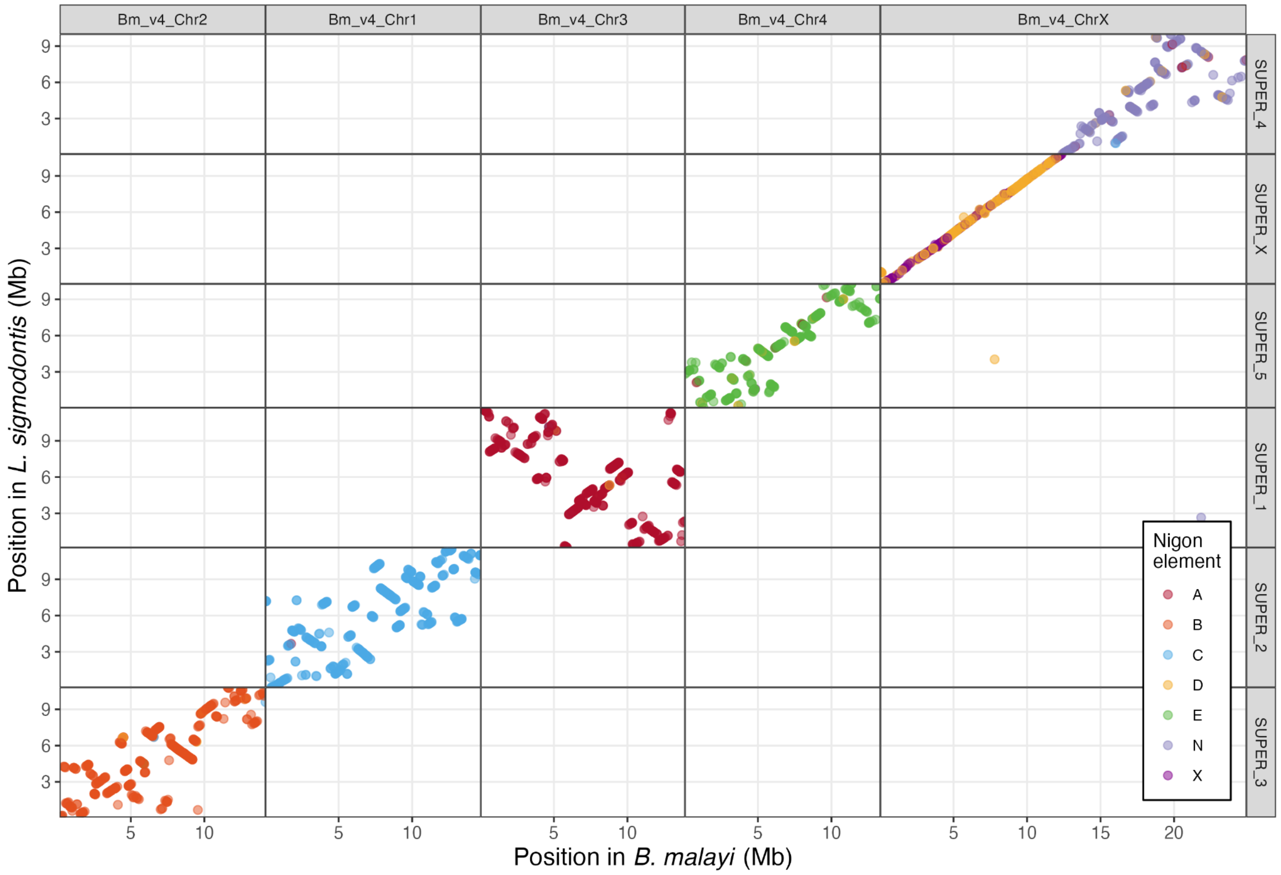


**Figure S5: Synteny between *B. malayi* and *L. sigmodontis***

The relative position of 2,209 BUSCO genes in the *B. malayi* and *L. sigmodontis* genomes. BUSCO genes are coloured by their Nigon assignment. Unassigned BUSCO genes are not shown. *B. malayi* chromosomes are ordered as in Figure 2B.

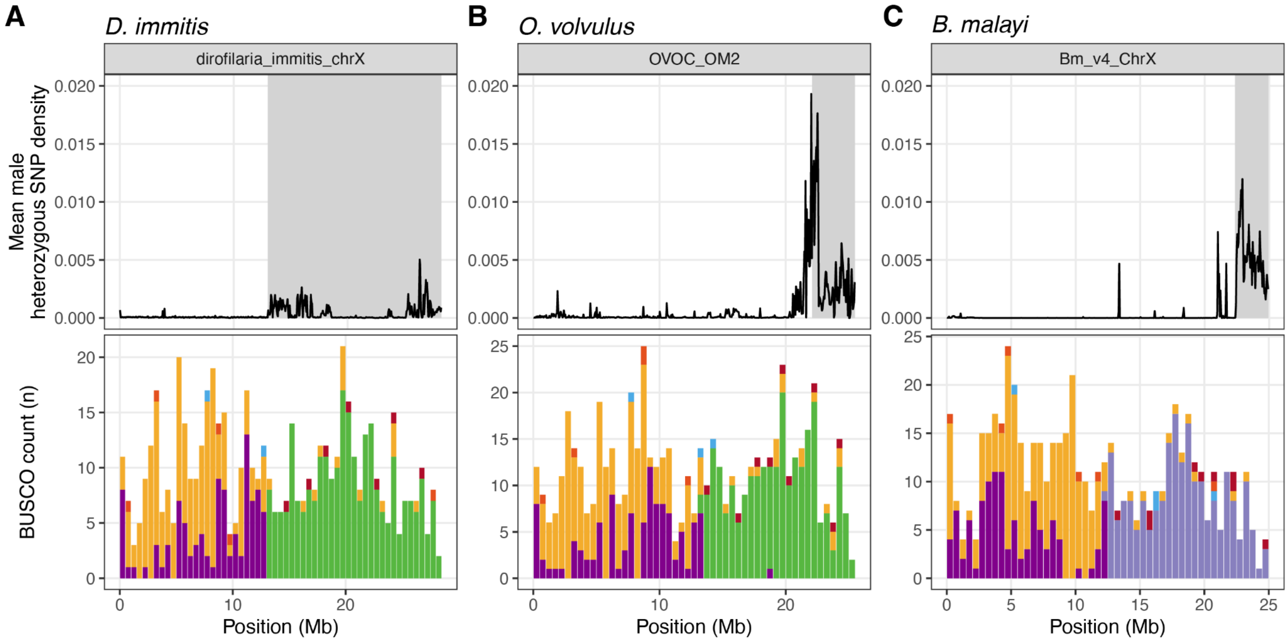


**Figure S6: Divergence between filarial X and Y chromosomes**

Mean male SNP density and Nigon element partitions in the X chromosomes of (A) *D. immitis*, (B) *O. volvulus*, and (C) *B. malayi*. Lines represent mean heterozygous SNP density in each 50 kb window using all male datasets for each species (Table S3). Pseudoautosomal regions are shown by grey shading. The histogram of locations of BUSCO loci allocated to Nigon elements (coloured as in Figure 1C) are binned in 500 kb windows.

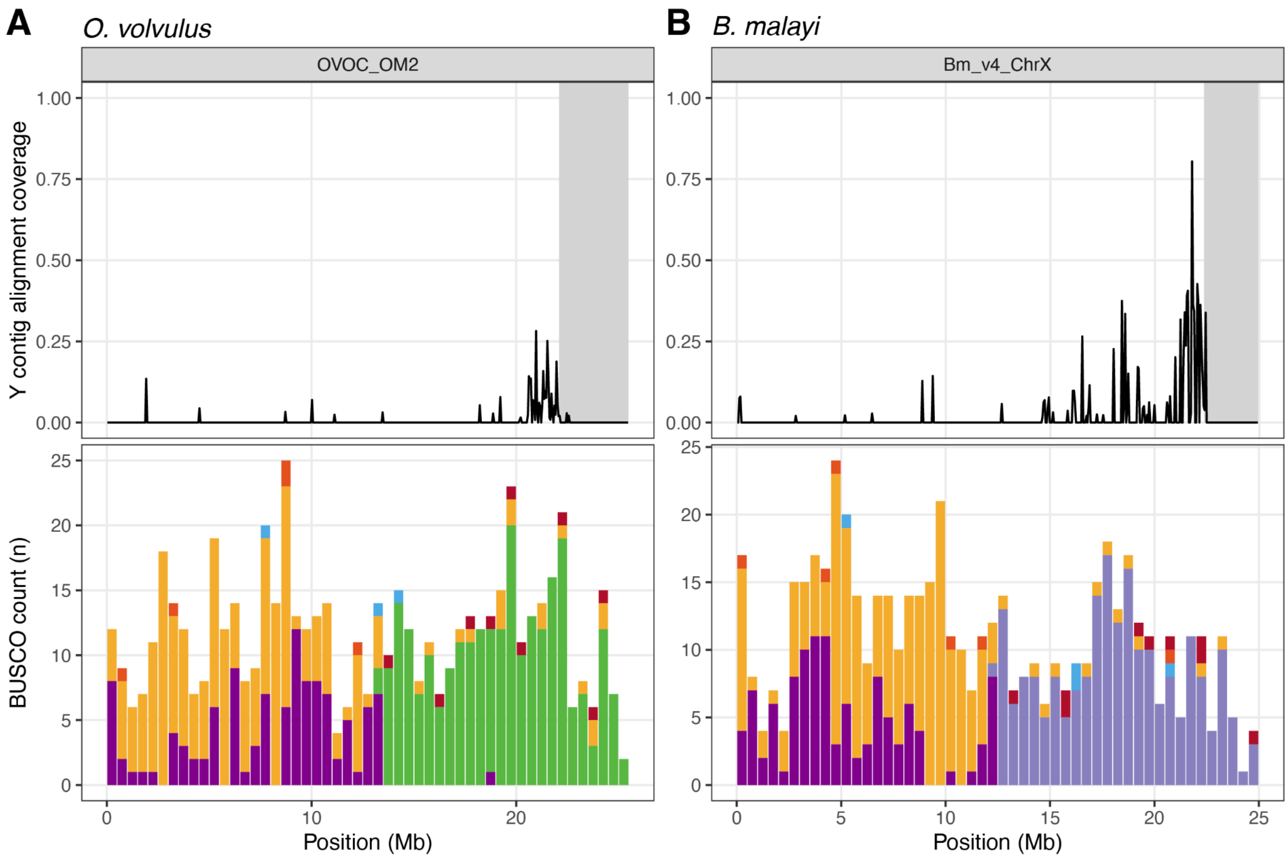


**Figure S7: Y contig alignment coverage in the *O. volvulus* and *B. malayi* X chromosomes**

Alignment coverage in 50 kb windows of the 148 *O. volvulus* Y contig identified by (Cotton et al. 2016) and 63 *B. malayi* Y contigs identified by (Foster et al. 2020) to the X chromosomes of *O. volvulus* and *B. malayi*. Only one-to-one alignments that were 1 kb or longer were considered. The histogram of locations of BUSCO loci allocated to Nigon elements (coloured as in Figure 1C) are binned in 500 kb windows.
